# Variability of Auditory Brainstem Responses in Diversity Outbred Mice

**DOI:** 10.1101/2024.04.21.590454

**Authors:** Roxana Taszus, Joaquin del Rio, Alexander Stoessel, Manuela Nowotny

## Abstract

Inbred mouse strains are mainly used in hearing research as animal models. Most of these lab mice strains consist of genetically identical individuals and natural variability as it is found in wild populations is not represented. Jackson Laboratories generated a new mice strain by an outbred of different mice strains to create mice with different pheno- and genotype, the Diversity Outbred strain (J:DO). By using 20 J:DO mice at the age of eight to nine weeks, we aimed to test the hearing variability in auditory thresholds and measured auditory brainstem responses at 15 different frequencies and sound pressure level from 0 to 80 dB SPL. Our measurements revealed a wide hearing bandwidth in J:DO mice that span over about 4.5 octaves. The high-frequency cut-off of the hearing threshold varied from 21 to 80 kHz and was on median at 64 kHz. Whereas the low frequency cut-off was at 3 kHz with low variation. The median characteristic frequency was found at 15 kHz and showed not a high variation in the sound level to induce an ABR threshold (20 dB SPL). We found not differences in hearing between male and female J:DOs. Future studies on hearing impairments in mice that aim to base on a genetically variable population could benefit from this new mouse strain and use our dataset on the hearing threshold in J:DO mice is a basis.

## 1. Introduction

Mice have become one of the universal model organisms in research in a wide range of fields, from medical study of pathologies to empirical analysis of still little understood mammalian abilities, like hearing and its responsible anatomy [e.g. 1,2,3]. Conventionally, inbred mouse strains have been preferred in hearing research. Their use mostly erases unpredictable genetic influences and permits the observation of experiments on elimination of chosen alleles connected to hearing ability [4,5,6], although genetic drift is still a factor known to cause some genetic heterogeneity [7].

In hearing research, typically a ton or click induced auditory response are used do observe minimal invasive hearing ability in mice, the auditory brainstem responses (ABR). The use of three electrodes allows the measuring of evoked potentials along the auditory ascending pathway, including the cochlea, auditory nerve, and brainstem nuclei [8,9]. In mice (*Mus musculus* Linnaeus, 1758), we expect to see up to five peaks in ABR recordings [10]. Depending on the placement of electrodes responses from: I) the auditory nerve; II) the cochlear nucleus, III) the superior olivary nucleus, IV) the lateral lemniscus, and V) the inferior colliculus can vary in their timing and amplitude response [11,12,13]. Hearing sensitivity in mice is subject to many factors, from genetic make-up [4], to type of anaesthesia [14], experiment procedure [15], and outside factors like noise pollution in housing [16]. However, we know from behavioural assessments of wild mice, that the hearing range of Mus musculus can range from 2.3 kHz up to 92 kHz [17].

We were interested in the variability of hearing capability in individuals and therefore decided to measure ABRs in a variable outbred strain. The J:DO mice allow for the assessment of a sample group approaching the genetic variability of human populations [18]. J:DOs are the outcome of continuous outbreeding of mice resulting from the Collaborative Cross to ensure a population with high degree of genetic and phenotypical variation [19,20,21]. The eight founder strains (A/J, C57BL/6J, 129S1/SvlmJ, NOD/ShiLtJ, NZO/HlLtJ, CAST/EiJ, PWK/PhJ, WSB/EiJ) are assumed to encapsulate about 90% of genetic variation in lab mice [22]. Most of these founder strains have previously been analysed and compared for a variety of factors, including hearing [e.g. 4], but not the J:DO strain. We have some insight from some founder strains into hearing thresholds, sensitivity and hearing loss. It has been shown that both A/J and NOD/ShiLtJ are effectively deaf at the age of 17-18 weeks (no response at click stimuli over 80 dB), and NZO/HlLtJ is at least hard of hearing (ABR responses at 75-80 dB), while WSB/EiJ and CAST/EiJ display higher extent of hearing sensitivity at that age at frequencies from 6 kHz to 30 kHz [23].

To date, ABR data of mouse strains were conducted at a variety of ages, with coarse resolution, and few frequencies, often just enough to establish general hearing ability or loss [4,24,25]. In our study we aimed to present a high resolution ABR audiogram of the J:DO mice, tested as young adults to investigate the range of individuality in hearing ability.

## 2. Materials and Methods

### 2.1. Animals

Jackson Laboratory Diversity Outbred (JDO, strain designation - JAX:009376) mice were used for measuring auditory brainstem responses. The sample consisted of 20 healthy individuals: 10 male and 10 female of different phenotypes (agouti n = 6:5, albino n = 2:2 and black n = 3:2, male:female). We tested the mice at the age of maturity at 8-9 weeks. The animals arrived at 4 weeks of age and were kept in open system cages in groups of 2-3 animals separated by sex. All animals had access to water and food (standard pellets) on demand, as well as nesting material, shelter, and enrichment (paper tubes and cotton balls). Humidity was kept at 55 ± 15%, artificial light provided 12-hour day and night cycles. Room temperature was adjusted to 20-24°C. The background noise level measured between 45-55 dB. All procedures were approved beforehand by the Administration Union for Animal Health and Foodstuffs Control Jena-Saale-Holzland under the designation FSU-10-002, thereby following the standards set by Directive 2010/63/EU on the protection of animals used for scientific purposes.

### 2.2. General Anaesthesia

During the preparation and experiment the animals were put under anaesthesia. We used a solution of ketamine (Ketavet, Pfizer GmbH, ca. 40-50 mg/kg) and medetomidine (Dormitor, Pfizer GmbH, ca. 0.40 – 0.50 mg/kg) for anaesthesia. After initial anaesthesia supplemental doses of 0.01 ml/h were administered by a narcotic pump (AL-1000 syringe pump, World Precision Instruments, Sarasota, USA).

The depth of anaesthesia was tested with the toe-pinch reflex in regular intervals outside, and via monitor inside the sound-attenuated experiment chamber. During the recording, ophthalmic ointment was applied to the eyes and the mice were placed on a heating pad monitored by a temperature control unit (HB 101, Panlab, Barcelona, Spain) to keep their body temperature at 37.5 - 38.5°C.

### 2.3. Auditory brainstem response measurements and statistical analysis

Hearing thresholds were obtained by playing pure-tone frequencies in increasing sound pressure levels from 0 to 80 dB SPL (step size 5 dB). Data recording, stimulus presentation and equipment control were organised by a MATLAB script (MATLAB 2022a, Natick, Massachusetts: The MathWorks Inc.) developed in-house. Three silver electrodes were fixed subcutaneously above the vertex (+), behind the left bulla (-), and a grounding electrode in the upper back. We measured 15 different frequencies (2, 3, 4, 8, 15, 16, 17, 21, 25, 32, 45, 55, 64, 72, and 80 kHz). The frequencies were chosen to concur with frequencies used in other studies, with a high resolution around the reported low- and high-end cut-offs, as well as the established characteristic frequency (CF). Here CF is defined as the frequency requiring the least stimulus to gain a measurable response. The frequencies between the established ones were chosen to grant a more detailed picture of an individual’s hearing curve. ABRs were measured with a randomized order of frequencies starting in one frequency in 5 dB step intensity from 0-80 dB before moving to the next to prevent habituation effect. Low- and high-end cut-off frequencies were set at the last frequencies ABRs under 60 dB could be detected. The acoustic stimuli were generated by a soundcard (Fireface 400, RME) and amplified (Rotel RBS 708x) before being played to the left ear with 10 cm distance between the speaker (R2904/7000, Scan-Speak, Videbæk, Denmark) directly facing the pinna of the animal. Evoked potentials were amplified with a DAGAN EX1 Differential amplifier (DAGAN Cooperation, Minnesota, US) before being fed back into the soundcard (sampling rate 184 kHz) and analysed in Matlab. The amplifier filters were set to 300 (low-cut) and 3000 (high-cut) with a gain of 200. The ABRs were evoked by pure tone stimulation with 3 ms duration and rise/fall of 1 ms after a delay of 10 ms. Each dB increment was averaged 300 times. To test the general hearing ability a click was measured at 16 kHz with a duration of 0.005s and rise/fall of 0.0005s.

To visually verify the ABR threshold at each measured frequency, the wave pattern of the brainstem responses was rendered in ascending dB steps (5 dB-incremental levels from 0-80 dB). All results were analysed by three independent observers with experience in ABR measurements and were consistent. Statistical significance was evaluated with an unpaired Wilcoxon test in RStudio (Version 1.4.1106, RStudio PBC). The data was visualised in R using the ggplot package (Villanueva and Chen, 2019), and SigmaPlot 15.0 software (Systat Software, San Jose, CA, USA). Boxplots present the median, the full ranges, and interquartile ranges (IQR). Levels of significance are indicated as follows: p < 0.05 = *, p < 0.01 = **, p < 0.001 = ***.

## 3. Results

We measured the ABR thresholds of 20 JDO mice in response to pure-tone stimuli from 2 to 80 kHz (Fig. 1). Regarding the most sensitive hearing, we found the median CF at 15 kHz with a median threshold of 20 dB SPL (IQR: 20 to 25 kHz, N = 20, Fig. 1B). At this CF, the measured sound pressure level varied from 15 to 35 dB SPL in the J:DO mice. At the stimulus frequency of 25 kHz, there was a high variability in the measured sound pressure level to induce an ABR threshold wave. The threshold difference between individuals rang at this frequency over 60 dB (from 20 to 80 dB SPL, N = 20). Concerning the upper and lower hearing ranges (threshold pass the level of 60 dB SPL), the median low-frequency cut-off was detected at 3 kHz (IQR 2 to 3, N = 20) and the high-frequency cut-off at 64 kHz (IQR 45 to 72, N = 20). The ABR threshold curve span from the lower to the upper cut-off a median hearing bandwidth of about 4.5 octaves (IQR: 3.9 to 5.0) in J:DO mice. Due to the increasing interquartile ABR threshold values (Fig. 1B, dark grey areas), we observe an increasing individual difference especially in the high-frequency hearing of the J:DO mice. The high-frequency cut-off (at 60 dB SPL), varying from 21 kHz to 80 kHz (Fig. 1B, light grey area). In relation to sex-specific differences in the hearing thresholds, we detect in both male and female mice individuals with very sensitive hearing and those whose ABRs are limited to frequencies below 21 kHz (Fig. 2). Overall, we observed no significant difference between male and female individuals (p > 0.05).

**Fig. 1.**
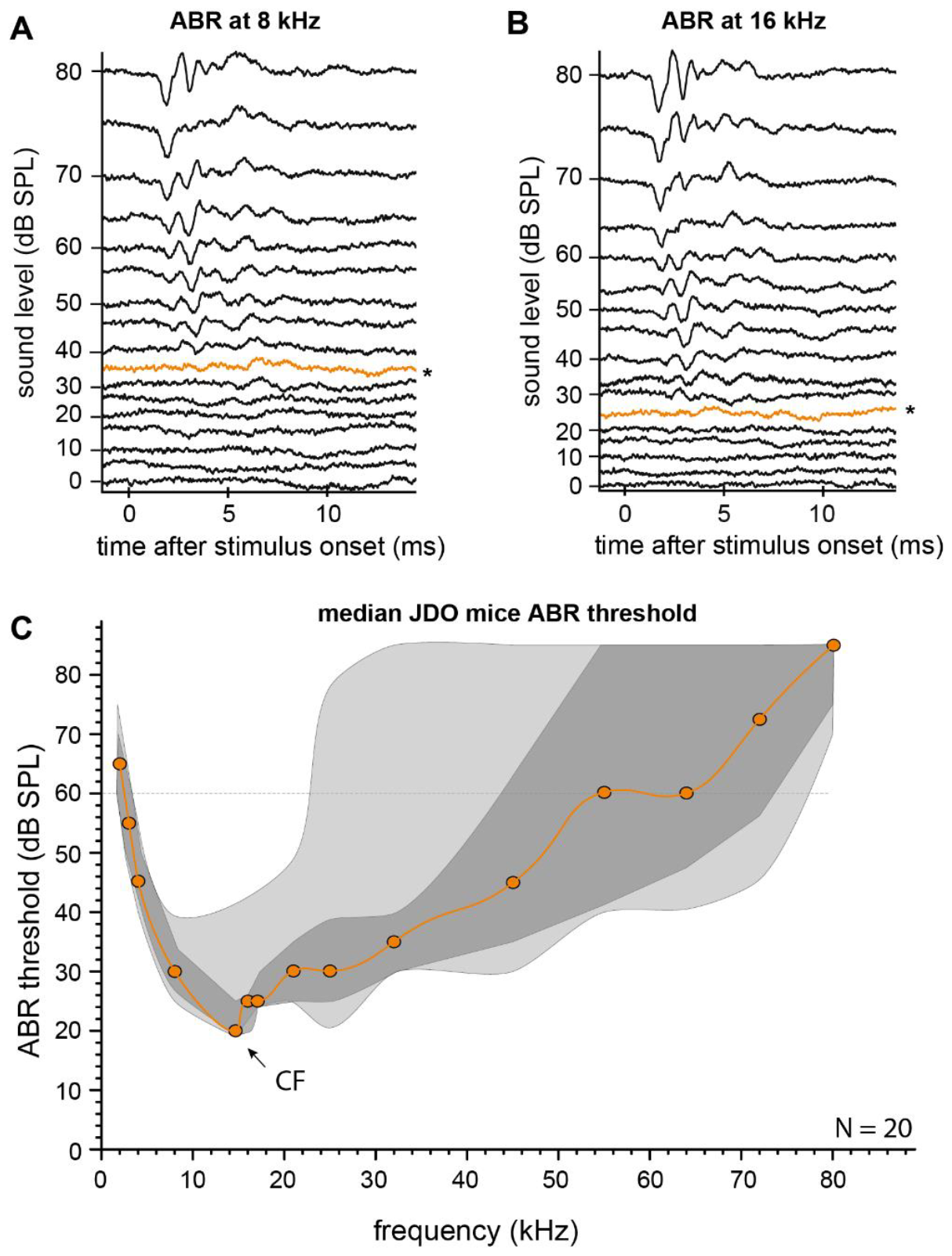
Auditory brainstem responses from JDO mice. A & B) Example of a pure-tone evoked ABR waveforms induced by a stimulus frequency of 8 (left panel) and 16 kHz (right panel) and increasing sound pressure level from 0 to 80 dB (5 dB steps). Hearing threshold is marked as * and an orange line. C) Median ABR thresholds for 20 JDOs with Q1 (25%) and Q3 (75%) area in dark grey and the total (5% and 95%) in light grey.

**Fig. 2.**
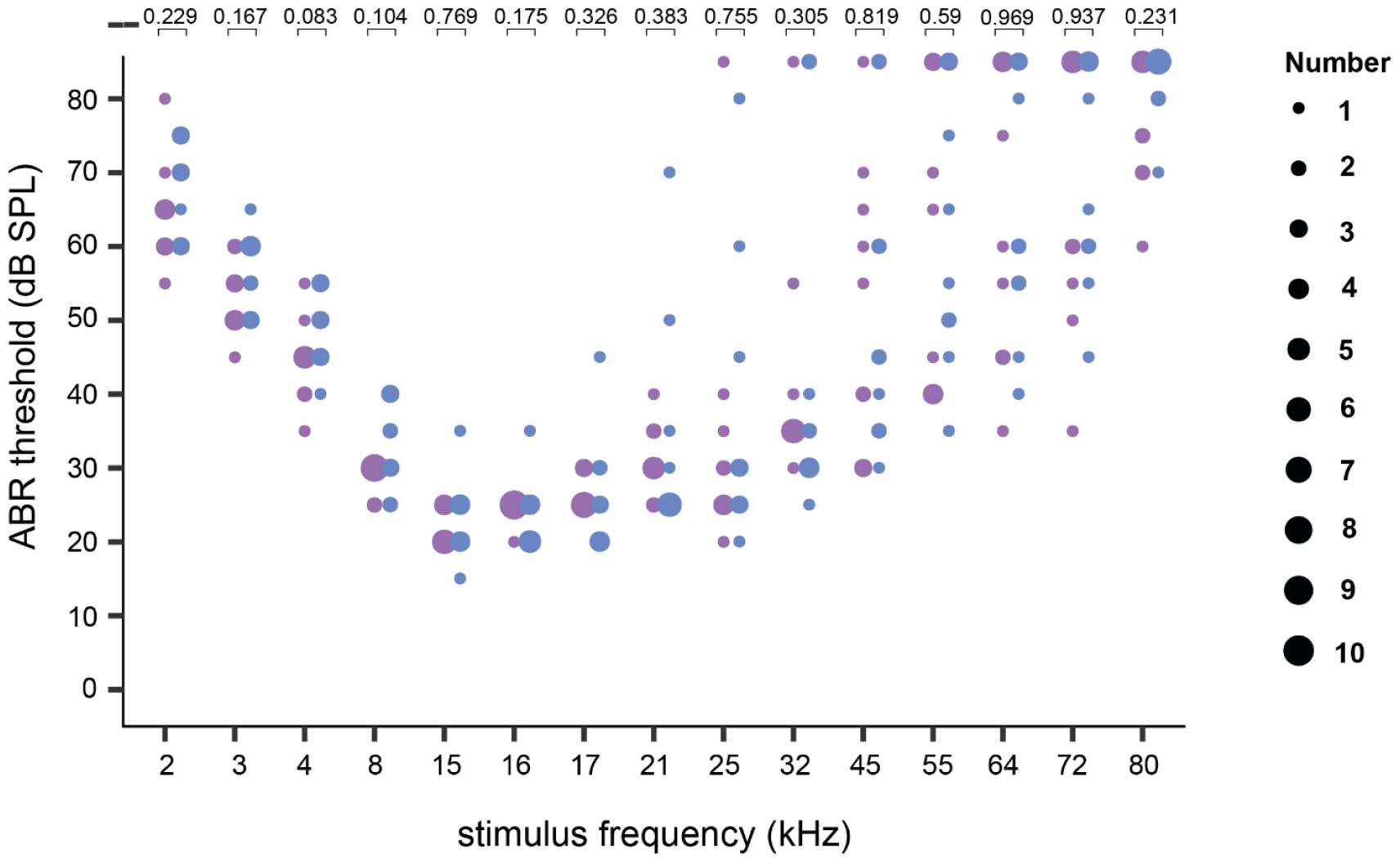
Distribution of ABR thresholds in male and female J:DO mice. ABR thresholds of male (blue, N = 10) and female (purple, N = 10) specimens plotted as bubble plots with statistical (Wilcoxon test) significant validation (p-value) of the threshold difference.

We further try to distinguish variances in hearing range for different phenotypes. At first, we have to mentioned that there is no possibility to order a fixed number of phenotypes from the breeder. We were able to distinguish three (i) agouti, ii) albino and iii) black) of the seven (black, albino, agouti, light bellied agouti, brown, tan, and black with blazes on their head) phenotypes that are mentioned by the breeder. The numbers of mice in our found phenotypes varied from 4 to 11 individuals (Fig. 3). These numbers lead to the consequence that we performed no statistic analysis between the phenotypes of J:DO mice. However, we found only minor differences in hearing thresholds between the three observed phenotypes (Fig. 3). With 11 agouti mice, we have the highest number for this phenotype. In this agouti group we found low variability in the low-frequency hearing and the CF. The high-frequency cut-off was highly variable (from about 21 to 77 kHz) in this phenotype (Fig. 3, left panel). With 35 dB SPL, an albino J:DO mice showed the lowest sensitive (highest hearing threshold) at 15 kHz with 35 dB SPL (Fig. 3, middle panel, dotted green arrow). On the other, hand we measured in an albino J:DO mouse a hearing range that had a high-frequency cut-off at 80 kHz and the highest hearing bandwidth at 60 dB SPL (Fig. 3, middle panel, solid green arrow). We notice that black J:DOs seemed to have overall lower and uniform ABR threshold values between 15 and 25 kHz (Fig. 3, right panel, N = 5). Indeed, we measured the highest sensitivity (lowest hearing threshold) in a black-coat mouse (JDO_14_30M) at 15 dB SPL on a 15 kHz stimulation (Fig. 3, black arrow in the right panel).

**Fig. 3.**
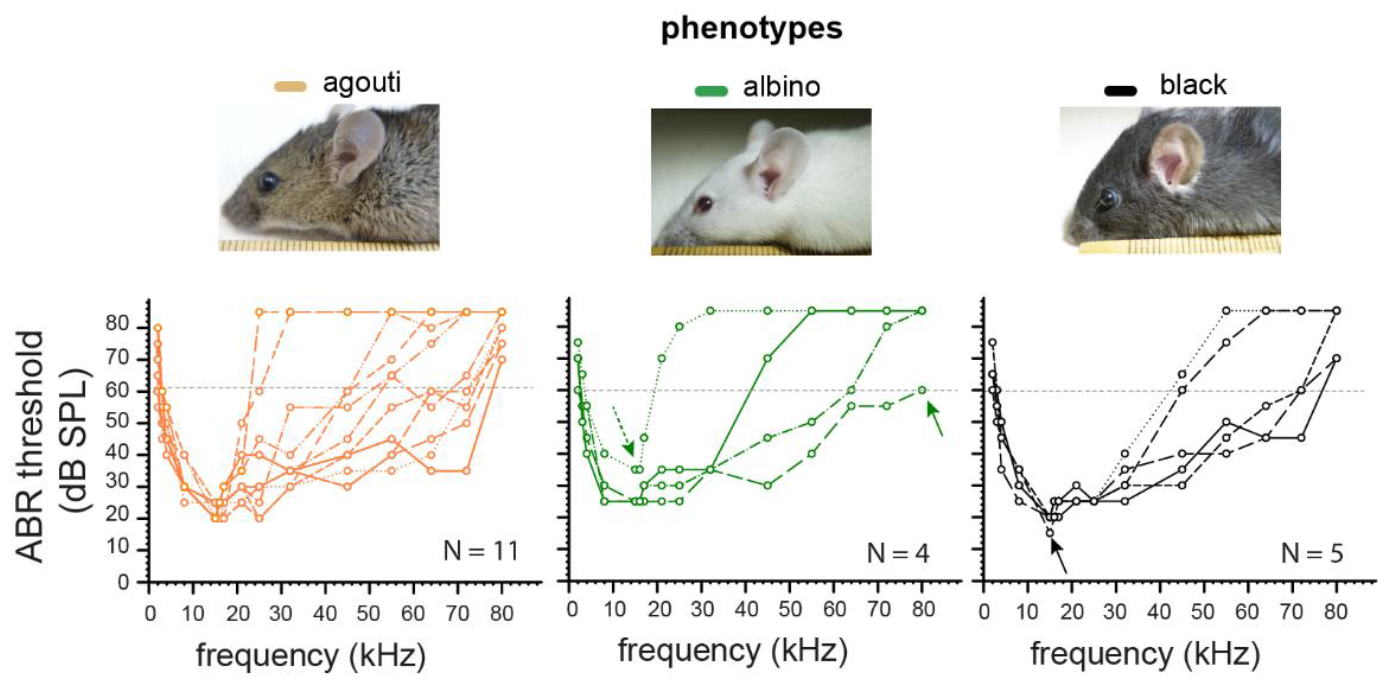
Phenotype-dependent ABR hearing thresholds for JDO individuals. Upper photographs depict the found phenotypes in our mice (agouti = orange, n = 11; albino = green, n = 4; black = black, n = 5). Below the corresponding individual ABR threshold are shown grouped by each phenotype. The grey dotted line at the ABR threshold of 60 dB SPL mark the hearing bandwidth.

## 4. Discussion

In order to investigate whether variability in hearing thresholds of J:DO mice reflects their reported genetic diversity [20,21,26,27,28], we measured ABRs of 20 individuals, 10 of each sex. We found for the J:DO strain a hearing capability that spans over a wide range of frequencies, from 3 to 80 kHz, with a median CF at 15 kHz on about 25 dB SPL. This hearing sensitivity is moderate compared to other mouse strains (Fig. 4, 16). However, the variability of the CF is rather low in this diversity outbreed strain. The low-frequency cut-off of the hearing range varied not much between the individuals and was at about 3 kHz. In contrast, the median higher-frequencies cut-off hearing thresholds, ranging between 21 and 80 kHz, was at median 64 kHz. This observation corresponds with finding by Du and colleagues that found their outbred CFW mice to fall into different patterns of hearing loss paralleled by patterns of ABR thresholds [29].

**Fig. 4.**
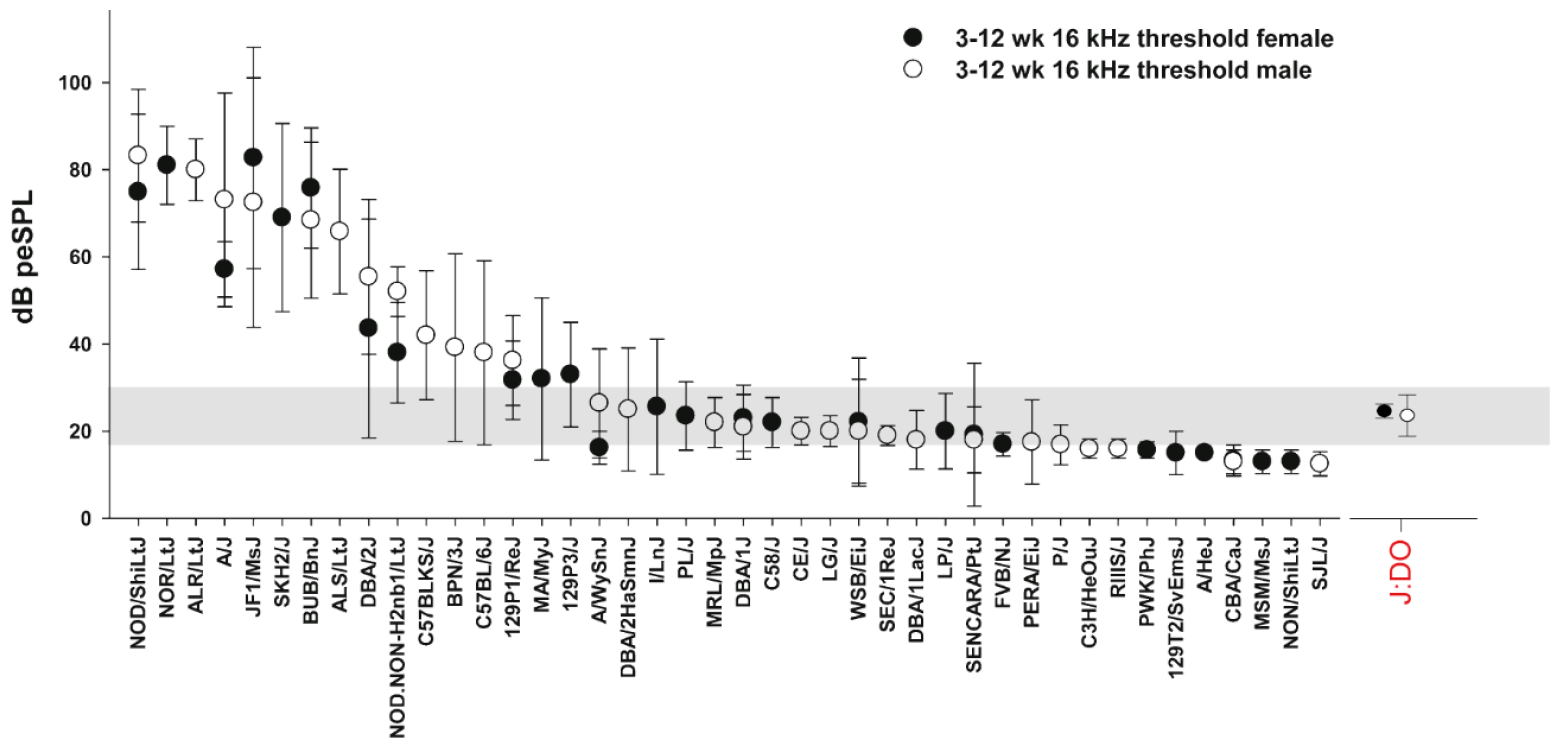
Allocation of the averaged ABR hearing thresholds of J:DO mice. Level-sorted ABR thresholds measured at 16 kHz for 42 inbred mouse strains with the new J:DO values at the right side. [changed after 17].

Our analysis on the differences in acoustic thresholds between male and female individuals shows no significant differences based on the sex for 8 to 9-weeks-old J:DO mice. Basic pre-clinical (meaning unaltered/not experimented on) analyses on possible hearing differences related to sex are still rarely reported for most lab mouse strains [30, 31,32]. However, there have been many investigations about the relationship between sex and hearing thresholds in mice in regards to age-related hearing loss and other noise-induced damage. Some of these studies present no differences in acoustic acuity between sexes in young pre-clinical mice [33,34,35] that align with our ABR measurements in the J:DOs, while other experiments suggest inherent variability linked to sex [36,37,38,39,40]. Whether there is a difference between males and females seems to depend on the mouse strain tested. We know that CBA/CaJ and CBA/J males have higher ABR thresholds than females in frequencies above 21 kHz at 50 days of age [32,36]. A comparable observation can be made in C57BL/6 mice, but only at > 100 days of age [37], at an age of 50 days the measurements of both sexes were nearly identical [41]. Young C57BL/6J mice tested at the age of 37 days display no significant difference between male and female [33]. In B6CBAF1/J mice, a cross between C57BL/6 and CBA/J, only the highest measured frequency, 32 kHz, shows a small but statistically significant lower threshold for female mice [42]. In summary it can be stated that although female mice of some strains have slightly lower thresholds at frequencies above 25 kHz, most differences in hearing thresholds between male and female individuals are a consequence of aging and hearing loss. Nevertheless, it has been shown that female mice are somewhat protected from hearing loss caused by trauma [e.g. 43], age [e.g. 37], and environmental factors like noise [e.g. 39,44] and diet [45], which was not tested in our study.

The diversity in this mouse strain was reached by the cross breeding of different mice strains, which lead to different phenotypes. Of the seven possible phenotypes among Diversity Outbred mice of 2022 (black, albino, agouti, light bellied agouti, brown, tan, and black with blazes on their head), we were able to identify three different phenotypes: i) albino, ii) agouti and iii) black mice. It is not possible to order by phenotypes. Only sex and age are criteria for selection when the mice were ordered. To test all seven phenotypes and both sexes for this strain with ten animals for each phenotype and sex, a number of at least 140 mice must be calculated and even higher numbers to compensate for the variable numbers of phenotype in the delivery by the breeder. To analyse all phenotypes in detail is not the intention when using this mice strain for experiments. It is more on showing individual differences and variability. Nevertheless, we group our ABR data for the three identified phenotypes and observed a trend for more sensitive hearing in black coat mice between 15 and 25 kHz. Whereas we observed the highest degree of variability in the albino group which contains individuals with hearing comparable to agouti specimens, but also severe deficits in the higher frequencies. It has previously been shown that melanin plays a role in the prevention of age-related hearing loss [46,47,48]. Furthermore, the lack of melanocytes can be linked directly to poor hearing acuity in mutant mice [49]. Although the lack of melanin alone has not been proven to affect ABR thresholds [50] adverse effects have been measured in cochlear nerve compound action potentials in gerbils [51].

In our sample 4 out of 20 mice were albinos with no pigmentation in their fur or eyes. Albinism is a rare phenotype in nature and especially in humans, estimated to occur in 1 of 20,000 births [52] and viewed as a symptom of mutations in at least six different genes [53]. All types of albinism have been reported to be accompanied by sensory deficits, particularly in visual and acoustic acuity [53,54]. It is therefore difficult to view the artificial genetic diversity of J:DO mice compared to natural wildtype mice or even human populations from this perspective. However, since the audiogram data for wild house mice have been obtained from only three individuals [17], which showed slight variation, they cannot inform sufficiently on the variation in hearing acuity of a natural mice population. In order to draw a comparison between artificially diverse and naturally diverse samples with any statistical significance, an experiment akin to the one presented here would have to be done on wild *Mus musculus*.

## CRediT authorship contributions

Roxana Taszus: Formal analysis, Investigation, Data curation, Validation, Writing - Original Draft, Writing - Review & Editing

Joaquin del Rio: Formal analysis, Review & Editing

Alexander Stoessel: Resources, Review & Editing

Manuela Nowotny: Conceptualization, Methodology, Validation, Supervision, Funding Acquisition, Project administrations, Resources, Writing - Original Draft, Writing - Review & Editing

## Data availability

The data can be made available on request.

## Acknowledgements

This endeavour would not have been possible without the technical expertise of Steven Abendroth and his familiarity with both equipment, and programming. We would also like to extend our gratitude to Toni Wöhrl for his advice on analysis in R. Funding was provided by the DFG (NO 841/11-1).

